# SARS-CoV-2 surveillance in Norway rats (*Rattus norvegicus*) from Antwerp sewer system, Belgium

**DOI:** 10.1101/2021.03.06.433708

**Authors:** Valeria Carolina Colombo, Vincent Sluydts, Joachim Mariën, Bram Vanden Broecke, Natalie Van Houtte, Wannes Leirs, Lotte Jacobs, Arne Iserbyt, Marine Hubert, Leo Heyndrickx, Hanne Goris, Peter Delputte, Naomi De Roeck, Joris Elst, Robbert Boudewijns, Kevin K. Ariën, Herwig Leirs, Sophie Gryseels

## Abstract

**Background:** SARS-CoV-2 human-to-animal transmission can lead to the establishment of novel reservoirs and the evolution of new variants with the potential to start new outbreaks in humans.

**Aim:** We tested Norway rats inhabiting the sewer system of Antwerp, Belgium, for the presence of SARS-CoV-2 following a local COVID-19 epidemic peak. In addition, we discuss the use and interpretation of SARS-CoV-2 serological tests on non-human samples.

**Methods:** Between November and December 2020, Norway rat oral swabs, feces and tissues from the sewer system of Antwerp were collected to be tested by RT-qPCR for the presence of SARS-CoV-2. Serum samples were screened for the presence of anti-SARS-CoV-2 IgG antibodies using a Luminex microsphere immunoassay (MIA). Samples considered positive were then checked for neutralizing antibodies using a conventional viral neutralization test (cVNT).

**Results:** The serum of 35 rats was tested by MIA showing 3 potentially positive sera that were later shown to be negative by cVNT. All tissue samples of 39 rats analyzed tested negative for SARS-CoV-2 RNA.

**Conclusion:** This is the first study that evaluates SARS-CoV-2 infection in urban rats. We can conclude that the sample of 39 rats had never been infected with SARS-CoV-2. We show that diagnostic serology tests can give misleading results when applied on non-human samples. SARS-CoV-2 monitoring activities should continue due to the emergence of new variants prone to infect Muridae rodents.

## Introduction

Emerging infectious diseases have been in the spotlight of scientific research in recent years. Most studies have focused mainly on the role of domestic and wild animals as zoonotic virus reservoirs and the phenomena that drive animal-to-human transmission in order to explain outbreak processes and spillover dynamics (e.g. Karesh et al 2010, Han et al 2016, Wardeh et al 2020). However, the possibility of human-to-animal viral transmission raised concern during the SARS-CoV-2 pandemic in 2020, when an asymptomatic dog from Hong Kong, whose owner was a COVID-19 patient, tested positive for the virus (Sit et al 2020). Since then, similar human-to-animal transmission events have been reported worldwide in domestic dogs (Sit et al 2020), cats (Chen at al 2020; Garligliani et al 2020), farmed minks (ECDC 2020, Oreshkova et al 2020, Hammer et al 2021, Oude Munnink et al 2021), and numerous zoo animals (McAloose et al 2020; OIE 2021). These events stimulated the scientific and public health community to better understand the implications and origins of this phenomenon.

The probability of human-borne SARS-CoV-2 emerging in animal populations differs between animal species through genetic and ecological differences (Gryseels et al., 2020). Susceptibility firstly depends on the ability of SARS-CoV-2 to enter host cells, which is determined by the affinity between the SARS-CoV-2 Receptor-Binding Domain (RBD) in the spike (S) protein and its binding receptor in host cells, Angiotensin-converting enzyme II (ACE2) protein (Othman et al 2020; Qiu et al 2020; Wu et al 2020). Whether the virus, after entering a host cell, can be transmitted persistently depends on individual characteristics, infection dynamics and ecological characteristics of the population. The longer the virus is shed from infected animals and / or the higher the contact frequency between animals, the likelier it can initiate a successful transmission chain. A good example of an optimal situation can be found in mink fur farms, which present a highly susceptible species (American mink *Neovison vison*) housed indoors in extreme high densities; leading to SARS-CoV-2 outbreaks as reported worldwide (ECDC 2020; Hammer et al 2021; Oude Munnink et al 2021). In nature, some mammals may also live in such high-density settings, particularly gregarious bats and fast-reproducing rodents. House mice (*Mus musculus*), Norway or brown rat (*Rattus norvegicus*) and the black or roof rat (*Rattus rattus*) are among the most ubiquitous rodents in the world (Feng & Himsworth 2014). They are considered true commensals, often living in close proximity to humans, increasing the risk of pathogen transmission, as they are a source of a wide range of viral, bacterial and parasitic zoonoses (Himsworth et al 2013). In Europe, Norway rats are well adapted to a synanthropic lifestyle and thrive in urban environments, including city sewer systems, where they find food, water and shelter (Mughini Gras et al 2012, Pascual et al 2020). Considering that many studies have detected SARS-CoV-2 in wastewater from the sewage system globally (e.g. Medema et al 2020; Randazzo et al 2020; Wu et al 2020), as well as in Antwerp, Belgium (Boogaerts et al 2021), these below-ground rodent populations can be exposed to SARS-CoV-2.

To date, only non-zoonotic *Betacoronaviru*ses were detected in Norway rats like Rat Coronavirus (RCov), China Rattus coronavirus HKU24 (ChRCoV HKU24) and Longquan Rl rat coronavirus (LRLV) (Decaro & Lorusso 2020), though some human endemic coronaviruses (OC43 and NL63) may have originated from a rodent reservoir (Corman et al 2018). SARS-CoV-2 has been shown to efficiently infect and replicate in Cricetid rodent species like the golden Syrian hamster, *Mesocricetus auratus* (Boudewijns et al 2020, Chan et al 2020, Sia et al 2020), the deer mouse (*Peromyscus maniculatus*) and the bushy-tailed woodrat (*Neotoma cinerea*) (Bosco-Lauth et al 2021). However, rodent species of the Muridae family, like house mice (*Mus musculus*) (Bosco-Lauth et al 2021) and Norway rats (Cohen 2020), were found not susceptible to infection by the ‘wild-type’ Wuhan SARS-CoV-2 strain. Their ACE2 receptor does not bind to this strain’s spike RBD *in vitro*. However, after serial passage in laboratory mice, SARS-CoV-2 evolves the ability to replicate efficiently in this host, thanks to a single substitution in the RBD, i.e. N501Y (Gu et al 2020). Remarkably, N501Y substitution has arisen repeatedly in SARS-CoV-2 lineages circulating in humans, most notable the variants of concern like B. 1.1.7, B.1.351 and P.1 (Yao et al 2021). This suggests that 1) SARS-CoV-2 can evolve relatively easily to infect a previous resistant species, and 2) several SARS-CoV-2 variants currently circulating have the inherent ability to infect *M. musculus* and potentially other species of the Muridae family.

For these reasons, in the present study we tested Norway rats inhabiting the sewer system of Antwerp, Belgium, for the presence of SARS-CoV-2 in November and December 2020, following a local COVID-19 epidemic peak by viruses mostly not carrying the N501Y substitution. In addition, we discuss the use and interpretation of SARS-CoV-2 serological tests on non-human samples.

## Materials and methods

### Study area

The study was conducted in the sewage system of the city of Antwerp (the Ruien) (51°13 ‘16.6 “N 4°23 ‘50.2 “E), Belgium, for 2 weeks during November - December 2020. The Ruien is an old network of small-scale waterways covered in 1882 that nowadays receives and directs the wastewater and the rainwater of the city of Antwerp to a water treatment plant (Marine & De Meulder 2016).

### Data collection

To test for the presence of SARS-CoV-2 in the sewage water at the exact location where Norway rats were trapped; eight water samples of 150 mL each were taken from flowing household sewage water in open parts of the sewage pipes on two different days during the rat trapping sessions. Samples were stored in individual tubes at 4°C and processed the next day. Up to 30 rat-live-traps baited with fish boilies (Decathlon – ‘taste’) were set out and checked every morning during 2 weeks; trapped rats were transported to a BSL-2^+^ laboratory at the Central Animal Facility, Campus Drie Eiken, University of Antwerp. Rats were euthanized with an overdose of isoflurane, and then weighed, measured and data of their species, sex and reproductive status were registered. Blood samples were collected in tubes without anticoagulant; serum was separated and stored at −20°C. Tissue samples of the kidney, lung, liver, and a 5 mm piece of colon were stored at −80°C. Oral swabs in PBS and feces samples in RNA later were also collected and stored at −80°C. All procedures were carried out under the approval of the University of Antwerp Ethical Committee for Animal Experiments (ECD code 2020-21).

### SARS-CoV-2 RNA and antibody detection

### Detection of SARS-CoV-2

*RNA in wastewater* Detection of SARS-CoV-2 RNA in the wastewater samples was done essentially as described in Boogaerts et al. (2021). Wastewater was first centrifuged at 4625g for 30 minutes at 4 °C in an Eppendorf 5910R Centrifuge (Aarschot, BE). The supernatant (40mL) was transferred to Macrosep Advance Centrifugal devices with Omega Membrane (100 kDa; Pall, New York, US) for centrifugal concentration according to the manufacturer’s instructions, and the concentrate was standardize to 1,5 ml with UltraPureTM DEPC-Treated Water (ThermoFisher Scientific). RNA extraction was done with the automated Maxwell PureFood GMO and Authentication RNA extraction kit. In brief, 200 µL of the concentrate was added to 200 µL cetylrimethylammonium bromide buffer and 40 µL proteinase K and the total volume was incubated for 10 minutes at 56 °C. This mixture was transferred to the sample well together with 300 µL lysis buffer after which automated RNA extraction was started in the Maxwell® RSC Instrument (Promega). The final elution volume was 50 µL. Amplifications with qPCR were performed in duplicate in 20 µL reaction mixtures using a 2x SensiFAST™ Probe No-ROX One-Step kit following Boogaerts et al (2021). A six-point calibration curve with a concentration between 10^5^ and 10^0^ copies/μL was constructed in ultrapure DEPC-treated water for quantification of the different genes of interest. The EURM-019 reference standard for the construction of the calibration curve was obtained from the Joint Research Centre (JRC, European Commission). The lower limit of quantification (LLOQ) was defined as the concentration in the lowest point of the calibration curve and was 10^0^ copies/μL. The LLOQ of the N1, N2 and E qPCR corresponded with Ct-levels of 36.1, 36.4 and 36.6, respectively.

### Serology

To test SARS-CoV-2 exposure in sewer rats, serum samples were first screened for the presence of binding anti-SARS-CoV-2 IgG antibodies, using an in-house Luminex microsphere immunoassay (MIA) (Mariën et al 2021). The MIA is a high-throughput test that allows the simultaneous detection of binding antibodies against different antigens of the same pathogen, increasing significantly the specificity of the test. However, the prediction performance of this test depends on the possibility to correctly estimating cut-off values of the negative controls. Since serum samples from sewer rats captured before the SARS-CoV-2 outbreak were not available, we used as negative controls serum from rats (*n*=7) trapped in forest and parks from Antwerp, outside the sewer system, as we considered that they were less likely to be exposed to SARS-CoV-2. Also, naïve laboratory mice (*n*=8) samples were used as negative controls. Positive control sera (*n*=10) were obtained from laboratory ifnar^-/-^ mice inoculated with a recombinant live-attenuated yellow fever virus that expressed the spike unit of SARS-CoV-2 (Sanchez-Felipe et al 2020). The MIA was run with two different beads coated with the virus ‘nucleocapsid and spike antigens (Ayouba et al 2020). A biotin-labelled goat anti-mouse IgG Y-chain specific conjugate (Sigma, B7022, 1/300 dilution) was used for visualization of the primary antibodies. Samples were considered to be positive if crude median fluorescence intensity values (MFI) were higher than 3x standard deviation (SD) of the negative control samples for both antigen-coated bead sets. All samples that were considered to be positive on the MIA (*n*=7) were checked for neutralizing antibodies using a conventional viral neutralization test (cVNT) (Mariën et al 2021). We only considered a sample to be seropositive if antibodies were detected on both the MIA and the cVNT.

### PCR tissues

Viral RNA was extracted from 140 µL of oral swabs samples in PBS and from 1 cm^2^ of feces using the QIAamp Viral RNA mini kit (QIAGEN, Valencia, California, USA) and from 30 mg kidney, lung, liver and colon samples using the NucleoSpin RNA mini kit (Macherey-Nagel, Düren, Germany) according to the manufacturer’s instructions. We tested for the presence of SARS-CoV-2 RNA via the CDC 2019-nCoV Real-Time RT-PCR protocol targeted to two regions of the nucleocapsid protein (N) gene, N1 and N2 (Lu et al 2020) performed on 5 µL of RNA using the SARS-CoV-2 (2019-nCoV) CDC RUO kit (IDT Cat. No. 10006713). The positive control used was a SARS-CoV-2 N synthetic probe (IDT, USA) designed for the present study. To monitor RNA extraction, we ran simultaneously a beta-actin (ACTB) assay as internal control (Borremans et al 2015) in a duplex assay N1/ACTB designed following Vogels et al (2020). N1/ACTB and N2 PCRs were performed separately for each sample with Luna Universal qPCR Master Mix (New England Biolabs) on an Applied Biosystems StepOne Real-Time PCR Instrument (Thermo Fisher Scientific) under the following thermal conditions: 52 °C for 10 min, 95 °C for 2 min, 44 cycles with 95 °C for 10 s and 55 °C for 30 s.

## Results

Of the 8 water samples tested, 4 samples had detectable Ct values for Sars-CoV-2. Two samples were positive below the LLOQ and two samples had Ct values that equaled with ± 7 gene copies per ml of wastewater.

Serum samples of 35 sewer rats were analyzed by MIA. Three had MFI values higher than the cut-off values of the negative controls for both nucleocapsid and spike IgG antibodies, but all remained lower than the MFI values of the positive control samples (Fig. 1). The three potentially positive sera and four other sera with high MFI values were subsequently checked for neutralizing antibodies by cVNT. All samples were seronegative for neutralizing antibodies; suggesting that the captured sewer rats had not experienced SARS-CoV-2 in their life.

**Figure 1:**
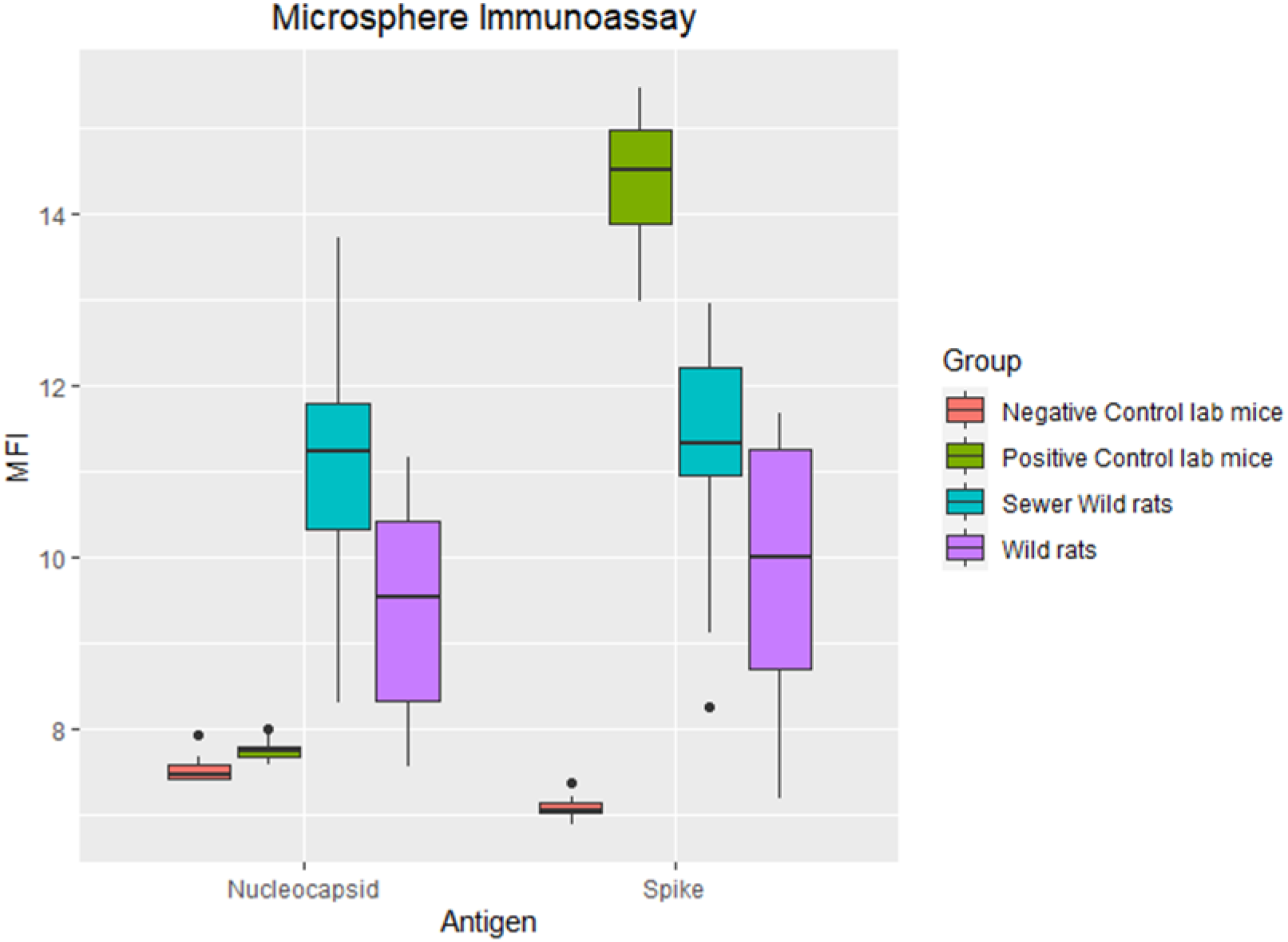
Boxplot showing the variation in log(MFI) values (Median Fluorescent intensities) for the different categories of mice/rats analysed in the microsphere immunoassay using the SARS-CoV-2 nucleocapsid and spike antigens.

Regarding the tissue samples analyzed, oral swabs, feces, colon, lung, liver and kidney samples of 39 sewer rats tested for the presence of SARS-CoV-2 by qRT-PCR were all considered negative.

## Discussion

To our knowledge, this is the first study that evaluates SARS-CoV-2 infection in urban Norway rats exposed to an environment contaminated with the virus, the sewer wastewater. According to the negative results obtained in both serology and PCR tests, we can conclude that the rodents studied had never been infected with SARS-CoV-2 despite continuous detection of viral RNA in the Antwerp sewer water (Boogaerts et al 2021), including sewer water collected at the exact location where the rats were captured.

Regarding the observed discrepancy between the results of the MIA and the VNT, we think it is worth mentioning that the interpretation of SARS-CoV-2 binding antibody tests (MIA or Elisa) should be made with care when used on different types of samples than what the assays were validated for. Indeed, although our MIA was clearly able to differentiate negative from positive control cases in laboratory mice (Fig. 1), it falsely categorized three wild type rats as positive when we estimated cut-off values based on serum from wild rats that were trapped outside of the sewers. The misclassification is explained by the fact that sewer rats had overall higher MFI values than rats trapped outside of the sewers (Fig. 1). This difference is likely caused by the higher exposure rate to many other pathogens in the sewer systems (dirtier conditions and higher population densities), which stimulates the adaptive immune system and results in overall higher binding antibody levels. Therefore, to confirm exposure to SARS-CoV-2 in a particular wildlife population based on serological data, VNTs are a better alternative (Tan et al 2020).

Studies to elucidate the animal species susceptible to SARS-CoV-2 have demonstrated the ability of the virus to spillover to several distantly related mammalian species (e.g. Chen at al 2020, McAloose et al 2020, Sit et al 2020, Hammer et al 2021), with the potential to stimulate the evolution of new variants with different antigenic properties (van Dorp et al 2020). This phenomenon can lead to various consequences, such as putting species conservation actions at risk if the virus affects endangered species, the establishment of novel reservoirs with the potential to start new outbreaks in humans, and the evolution of novel variants that may evade antibodies generated in humans, forcing the development of new antiviral therapies (Gryseels et al 2020, Mercatelli & Giorgi 2020, Hammer et al 2021, Oude Munnink et al 2021).

The absence of SARS-CoV-2 in our sample of Norway rats could possibly be explained by the dominance of SARS-CoV-2 lineages without the spike N501Y substitution in humans prior and at the time of sampling the rats. Since the beginning of the SARS-CoV-2 pandemic, many new variants have been involved in humans and in non-human animal hosts (van Dorp et al 2020; Hodcroft 2021; Leung et al 2021; Mercatelli & Giorgi 2020). Some of the currently most widespread variants, like B.1.1.7/501Y.V1, B.1.351/501Y.V2 and P.1/501Y.V3 that emerged from the UK, South Africa and Brazil, are potentially able to infect previous resistant species, such as Muridae rodents, thanks to the N501Y substitution in the RBD (Gu et al 2020, Yao et al 2021). This scenario, in conjunction with the synanthropic habits of several Muridae rodents and their ability to develop high-density populations, creates the ideal conditions for the spread of new epidemics. As such, despite the negative results found in Norway rats in the present study, we emphasize the need to carry out regular monitoring activities for the presence of SARS-CoV-2 in Muridae rodents, as well as other mammals exposed to humans, in order to detect human-to-animal transmission events and prevent future outbreaks emerging from new animal reservoirs.

## Acknowledgements

This work was funded through the 2018-2019 BiodivERsA joint call for research proposals, under the BiodivERsA3 ERA-Net COFUND programme (HL); FED-tWIN OMEgA (SG,HL); Research Foundation Flanders (FWO) (G0G4220N and G054820N to KKA) and the Research Council (BOF) of the University of Antwerp (to PD, project number: FFB200184). HL is member of the University of Antwerp Center of Excellence VAX-IDEA.

We are very grateful to Flinn De Vleeschauwer, water-link (in particular Werner Van Den Bogaert) and the Ruien operator Werkmmaat (in particular Willem Storms and Rasool Rezai) for enabling access to the trapping sites and their tremendous help during the trapping work.

